# Transcript assembly improves expression quantification of transposable elements in single cell RNA-seq data

**DOI:** 10.1101/2020.07.31.231027

**Authors:** Wanqing Shao, Ting Wang

## Abstract

Transposable elements (TEs) are an integral part of the host transcriptome. TE-containing noncoding RNAs (ncRNAs) exhibit considerable tissue specificity and play crucial roles during development, including stem cell maintenance and cell differentiation. Recent advances in single cell RNA-seq (scRNA-seq) revolutionized cell-type specific gene expression analysis. However, scRNA-seq quantification tools tailored for TEs are lacking, limiting our ability to dissect TE expression dynamics at single cell resolution. To address this issue, we established a TE expression quantification pipeline that is compatible with scRNA-seq data generated across multiple technology platforms. We constructed TE containing ncRNA references using bulk RNA-seq data and demonstrated that quantifying TE expression at the transcript level effectively reduces noise. As proof of principle, we applied this strategy to mouse embryonic stem cells and successfully captured the expression profile of endogenous retroviruses in single cells. We further expanded our analysis to scRNA-seq data from early stages of mouse embryogenesis. Our results illustrated the dynamic TE expression at pre-implantation stages and revealed 137 TE-containing ncRNA transcripts with substantial tissue specificity during gastrulation and early organogenesis.

## Introduction

Transposable elements (TEs) occupy a large proportion of eukaryotic genomes, representing roughly 50% of the human genome and 40% of the mouse genome. Though once regarded as non-functional parasitic sequences, mounting evidence suggests that TEs play pivotal roles in gene regulation. During evolution, TEs rewire host transcription networks through transposition, resulting in a wide variety of TE-derived regulatory elements, including promoters, enhancers, transcription terminators and chromatin loop anchors (Feschotte and Gilbert 2012; Rebollo et al. 2012; Cowley and Oakey 2013; Garcia-Perez et al. 2016; Chuong et al. 2017; Sundaram and Wysocka 2020). In the present-day, despite losing most of their transposition abilities, TEs continue to impact host genomes through transcription, which generates protein-coding-TE chimeric RNAs as well as noncoding RNAs (ncRNAs) that are crucial for normal and cancer development (Gifford et al. 2013; Hadjiargyrou and Delihas 2013; Hutchins and Pei 2015; Anwar et al. 2017; Rodriguez-Terrones and Torres-Padilla 2018).

TEs are major contributors of ncRNAs in both human and mouse. More than two thirds of mature long ncRNAs contain at least one TE and almost half of the total base pairs of long ncRNA are derived from TEs (Kelley and Rinn 2012; Kapusta et al. 2013; Veselovska et al. 2015). TE-containing ncRNAs show substantial developmental stage and tissue specificity and play essential roles during embryonic stem cell (ESC) maintenance and early embryogenesis. For instance, endogenous retroviruses (ERVs) are highly expressed in ESCs and are involved in ESC self-renewal and differentiation (Macfarlan et al. 2012; Santoni et al. 2012; Fort et al. 2014; Lu et al. 2014; Ohnuki et al. 2014; Wang et al. 2014). During mouse and human embryogenesis, a large number of TEs, including ERVs, long interspersed nuclear element-1 (LINE-1) and short interspersed elements (SINEs), become active and contribute to a significant proportion of total RNAs before blastocyst stage (Kigami et al. 2003; Peaston et al. 2004; Svoboda et al. 2004; Maksakova and Mager 2005; Fadloun et al. 2013; Göke et al. 2015; Grow et al. 2015; De Iaco et al. 2017; Ge 2017; Hendrickson et al. 2017; Jachowicz et al. 2017; Whiddon et al. 2017; Percharde et al. 2018). Moreover, knocking down specific TE families, including LINE-1 and MuERV-L, results in severe developmental defects (Kigami et al. 2003; Huang et al. 2017; Jachowicz et al. 2017; Percharde et al. 2018).

Despite the importance of TEs, quantifying TE expression using high-throughput sequencing data has been challenging. Due to TEs’ repetitive nature, sequencing reads that overlap with TEs are often discarded as a result of ambiguous mapping. Recently, several software tools were developed to address this issue and they enabled TE expression quantification in bulk RNA-seq data (Day et al. 2010; Jin et al. 2015; Srivastava et al. 2016; Lerat et al. 2017; Guffanti et al. 2018; Jeong et al. 2018; Valdebenito-Maturana and Riadi 2018; Bendall et al. 2019; Yang et al. 2019). In order to quantify the expression of repetitive elements, these tools either aggregate multi-aligned reads at TE subfamilies/families or redistribute them to individual TEs based on heuristic or statistical rules. Although proven to be successful in a range of biological systems, applications of the current TE quantification strategies were mostly limited to bulk RNA-seq, which lacks the ability to distinguish cell type specific TE expression.

Recent developments in single cell RNA-seq (scRNA-seq) provide unprecedented opportunities for examining cell type specific TE expression. However, effective TE quantification tools optimized for scRNA-seq data are lacking. Although the assessment of genome-wide transcriptional activity of TEs in single cells has been attempted by counting signals at TE subfamilies/families (Göke et al. 2015; Ge 2017; Boroviak et al. 2018; Brocks et al. 2018; Yandim and Karakülah 2019; He et al. 2020; Jonsson et al. 2020), such approaches are not optimal. Compared with bulk RNA-seq, scRNA-seq signal is much noisier and often shows 5’ or 3’ end enrichment along the transcripts. Counting reads at individual TEs or subfamilies/families fails to take into account the structures of the full-length transcripts, which can consist of multiple TEs from different subfamilies/families. Consequently, different expression values will be assigned to individual TEs within the same transcript. This caveat is especially obvious when dealing with scRNA-seq datasets where sequencing reads are enriched at either the 5’ or 3’ end of the RNA. Counting reads without the knowledge of the full-length transcripts will only capture TEs near the 5’ end or the polyA signal, resulting in an inaccurate picture of the genome-wide TE expression pattern.

In order to address these issues, we have developed an analytical pipeline tailored for TE expression quantification in scRNA-seq datasets. By comparing TE derived RNA-seq reads in bulk and scRNA-seq datasets, we first show that higher percentages of reads are mapped to TEs in scRNA-seq regardless of library preparation protocols. We further demonstrate that counting scRNA-seq signal at individual TEs leads to large amounts of false positives and generate bias due to 3’ enrichment of scRNA-seq signal. To overcome these challenges, we quantify TE expression in single cells using TE transcripts assembled from bulk RNA-seq. This approach successfully captures TEs that are exonized into ncRNAs and significantly improves scRNA-seq analysis by enriching for regions with true signal. Furthermore, applying our strategy to mouse early embryogenesis captured dynamic TE expression during pre-implantation stages. Expanding our analysis to mouse gastrulation and early organogenesis revealed 137 TE transcripts with substantial tissue specificity. These TE transcripts are mostly un-annotated and transcripts with different tissue enrichment show distinct TE composition. In summary, our study provides a systematic evaluation of TE derived reads in scRNA-seq datasets and establishes an effective computational approach for quantifying TE expression using scRNA-seq data.

## Results

### A higher percentage of reads are mapped to TEs in scRNA-seq compared with bulk RNA-seq

Due to the biological significance of TE-containing ncRNAs, we decided to focus our analysis on TEs that are not part of protein-coding transcripts. First, to determine the fraction of reads that can be mapped to these TEs in scRNA-seq data, we processed 36 publicly available single cell datasets (Supplemental Table S1). These datasets contain both human and mouse samples and were generated using 7 different scRNA-seq protocols. Bulk RNA-seq datasets from the same study or derived from the same cell line were included as controls. To preserve reads that originate from repetitive regions, multiple mapping was implemented during the alignment step. Reads that mapped to multiple locations or overlapped with more than one feature were distributed equally for signal quantification. Calculating the number of mappable reads based on genomic locations revealed that a large proportion of reads overlap with TEs in all tested scRNA-seq datasets, suggesting that TE expression can be captured by scRNA-seq (Fig. 1A). Notably, we observed a higher percentage of reads mapped to TEs in scRNA-seq compared with bulk RNA-seq, a phenomenon that was consistent across different scRNA-seq platforms even when only uniquely mapped reads were considered (Fig. 1A, Supplemental Fig. S1A).

**Figure 1.**
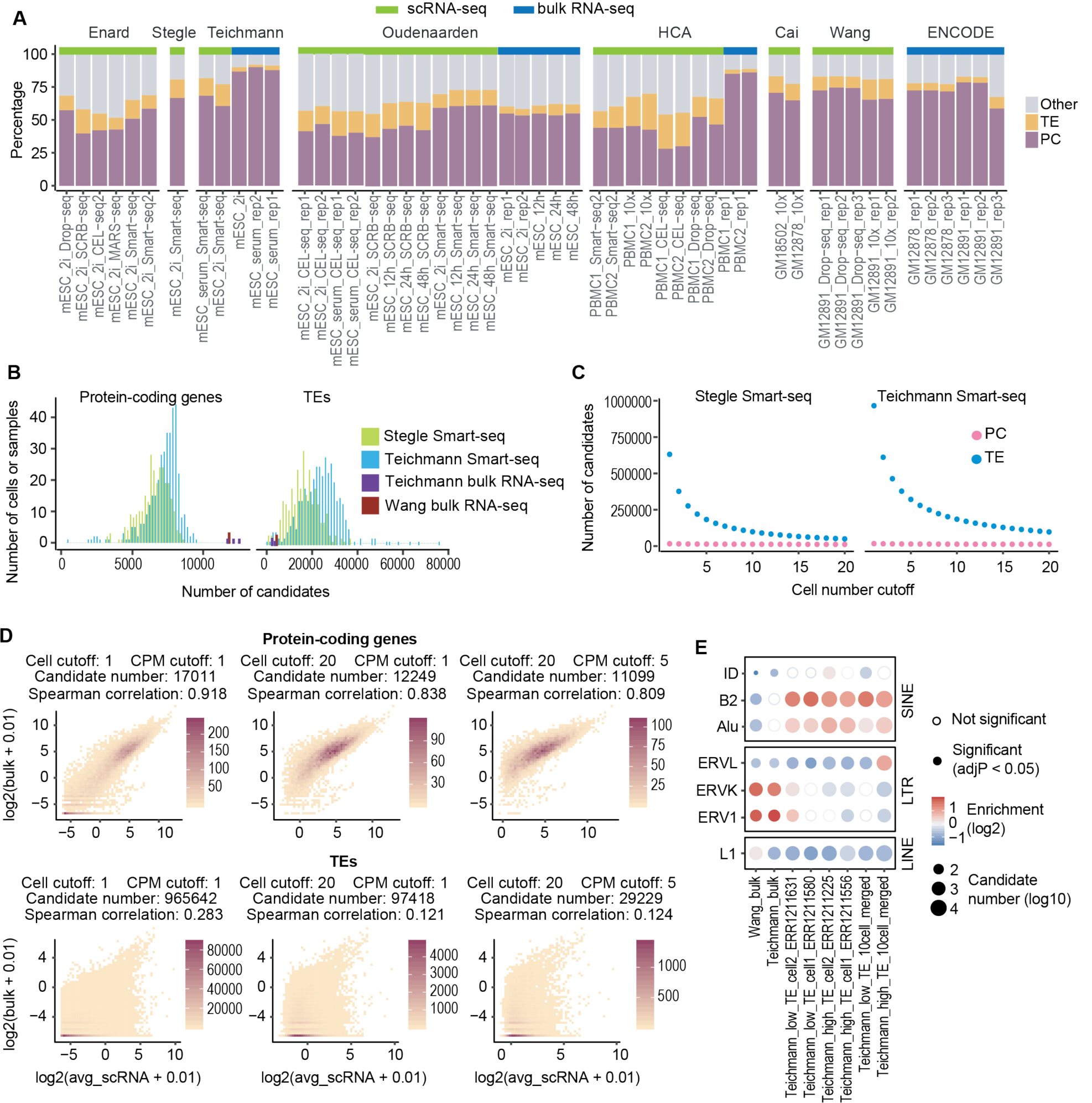
Counting scRNA-seq signal at individual TEs results in large numbers of false positive candidates. (A) Distribution of mappable reads in 16 bulk RNA-seq and 36 scRNA-seq datasets. Compared to bulk RNA-seq, scRNA-seq data have higher percentage of reads mapped to TEs. Samples were arranged by studies. PC: protein-coding exons defined by refSeq. TE: transposable elements that do not overlap with protein-coding exons. Other: other genomic locations. mESC: mouse embryonic stem cell. PBMC: human peripheral blood mononuclear cell. GM12878 and GM12891: human lymphoblastoid cell lines. (B) Number of expressed (Counts Per Million, CPM >=1) protein-coding genes and TEs in mESC bulk RNA-seq and Smart-seq samples. On average, 12,000 protein-coding genes and 6,000 TEs were detected in each bulk RNA-seq sample. In contrast, scRNA-seq captured 7,000 protein-coding genes and 20,000 TEs per cell. (C) Number of candidates as a function of cell number cutoff (the minimum number of cells each candidate is expressed in. Expression cutoff: CMP >=1). Although the majority of protein-coding gene candidates were consistently detected in mESC Smart-seq data, a large number of TE candidates were detected in less than 10 cells. (D) Correlation between bulk RNA-seq and averaged scRNA-seq signal at protein-coding genes and TEs (Teichmann lab, mESC). Low correlation between bulk RNA-seq and averaged Smart-seq signal was observed at TEs regardless of expression cutoff. Cell cutoff: the minimum number of cells each candidate is expressed in. CPM cutoff: the minimum CPM value for one candidate to be considered as expressed. (E) TE-family enrichment analysis using TE candidates identified from mESC bulk RNA-seq and Smart-seq. Enrichment of ERV elements was observed with bulk RNA-seq data, but not in single cells. Smart-seq data of four single cells with different percentage of TE reads and merged Smart-seq data from 10 cells were included.

To evaluate whether the high TE mapping percentage in scRNA-seq was associated with data quality per cell, we further compared scRNA-seq data generated using Smart-seq, 10x Genomics Chromium, SCRB-seq and Drop-seq and examined the relationships between the percentage of TE reads per cell and the two key parameters indicative of scRNA-seq quality: sequencing depth and the percentage of mitochondrial reads (Ilicic et al. 2016). Our analysis revealed that a high TE mapping percentage was observed across individual cells with limited correlation to sequencing depth (Supplemental Fig. S1B, C). Similarly, no strong correlation was detected between the percentage of TE reads and that of the mitochondrial reads (Supplemental Fig. S1C). These results suggest that the high TE mapping ratio in scRNA-seq is unlikely an artifact caused by extreme sequencing depth or cell death.

To address the concern that genomic DNA contamination may contribute to the majority of TE signal in scRNA-seq, we next quantified the number of total reads mapped to 5 non-overlapping genomic regions: protein-coding exons, TEs within the introns of protein-coding genes, other intronic regions of protein-coding genes, intergenic TEs, and other intergenic regions. A higher percentage of total reads were mapped to the intronic regions of protein-coding genes in scRNA-seq and the majority of TE overlapping reads were located within introns, suggesting that the signal originates from un-processed RNAs (Supplemental Fig. S1A). Furthermore, consistent with a previous report (La Manno et al. 2018), we observed that regions enriched for un-spliced scRNA-seq reads tend to be flanked by AT-rich sequences (Supplemental Fig. S1D). Taken together, these observations suggest that scRNA-seq TE signals are unlikely to come from genomic DNA contamination but rather are the products of the polyA priming step during cDNA synthesis.

### Counting scRNA-seq reads at individual TEs leads to large numbers of false positive candidates

Current TE expression analyses often quantify RNA-seq signal at individual TEs or TE subfamilies/families (Day et al. 2010; Jin et al. 2015; Lerat et al. 2017; Jeong et al. 2018; Yang et al. 2019; He et al. 2020; Jonsson et al. 2020). Our observation that a large proportion of scRNA-seq reads map to TEs, especially intronic TEs, raises the concern that counting reads at single TEs or TE subfamilies/families will aggregate noise and fail to exclude TEs that are part of protein-coding genes, resulting in high numbers of false positive candidates. To test this, we applied a similar strategy and analyzed bulk and Smart-seq datasets generated using mouse embryonic stem cells (mESCs) cultured in 2i medium (Buettner et al. 2015; Kolodziejczyk et al. 2015). Because mESCs represent a relatively homogeneous population and reads generated by Smart-seq and bulk RNA-seq share a similar distribution along the gene body (Ramsköld et al. 2012), we expected that the expression profiles obtained with scRNA-seq to be similar to those with bulk RNA-seq.

We first calculated the number of expressed protein-coding genes and TEs (CPM >=1) in these datasets. On average, 12,000 protein-coding genes and 6,000 TEs were detected in bulk RNA-seq samples. In contrast, scRNA-seq captured an average of 7,000 protein-coding genes and 20,000 TEs per cell (Fig. 1B). To evaluate the quality of these TE candidates, we examined the following three parameters: 1) The number of cells each candidate is expressed in, 2) the correlation between the signal in bulk RNA-seq and the average signal across single cells, 3) the over-represented TE families among all candidates. We reasoned that a true candidate should be expressed in a relatively large number of mESCs and exhibit a strong correlation between its bulk RNA-seq and averaged scRNA-seq signal. Strikingly, only protein-coding genes matched this expectation (Fig. 1C, D, Supplemental Fig. S2A). A large proportion of TE candidates were only detected in a small number of cells and exhibited weak correlations between scRNA-seq and bulk RNA-seq signal regardless of the expression cutoff. This observation remained valid after we performed the same analysis by counting signals from individual exons. The exon length distributions were comparable to those of TEs, ruling out the possibility that length discrepancy between TEs and protein-coding genes contributes to false positive TE candidates (Supplemental Fig. S2B-D). We further compared over-represented TEs within candidates identified from bulk RNA-seq and scRNA-seq by performing a TE-family enrichment analysis (Supplemental Fig. S3A). Although ERV elements have been shown to be expressed in stem cells (Macfarlan et al. 2012; Santoni et al. 2012; Fort et al. 2014; Lu et al. 2014; Ohnuki et al. 2014; Wang et al. 2014), they were only enriched in bulk RNA-seq in this analysis (Fig. 1E). scRNA-seq candidates obtained from this analysis were depleted of ERVs and instead enriched for SINEs (Fig. 1E, Supplemental Fig. S3B), which are often found near protein-coding genes and provide sequences that could act as reverse transcription priming sites (Medstrand et al. 2002).

In summary, the extreme high number of TE candidates obtained from scRNA-seq, the weak signal correlation between individual cells, as well as the discordance between bulk and scRNA-seq strongly suggest that counting scRNA-seq reads at individual TEs will result in large numbers of false positive candidates.

### Transcript assembly improves TE expression analysis

Transcript annotation serves as the cornerstone for expression quantification. Our ability to accurately assess the expression of protein-coding genes relies on well-annotated gene structures, which help to focus analysis on genomics regions with true signal. Although individual TEs are well annotated, it is usually unclear which TEs are expressed in a biological system and what the underlying transcript structures are. We reason that the large number of false positive candidates in scRNA-seq analysis is due to counting sparse and noisy signal at millions of TE copies, of which only a small proportion are truly expressed (Supplemental Fig. S3C). Therefore, we hypothesize that incorporating transcript structures of TE-containing ncRNAs into the analysis should help to reduce noise.

To obtain ncRNAs with exonized TEs, we performed transcript assembly using mESC bulk RNA-seq data. We identified 692 transcripts that overlap with TEs but not the exons of protein-coding genes (Fig. 2A, B, Supplemental Fig. S4). These transcripts were termed TE transcripts. To test the accuracy of our assembly, we focused on the promoters of assembled TE transcripts and examined several genomic signatures that are indicative of active transcription. Indeed, the majority of our TE transcript promoters overlap with FANTOM5 CAGE peaks (FANTOM Consortium and the RIKEN PMI and CLST (DGT) et al. 2014) and are enriched for ATAC-seq signal while depleted of CpG methylation (Fig. 2C).

**Figure 2.**
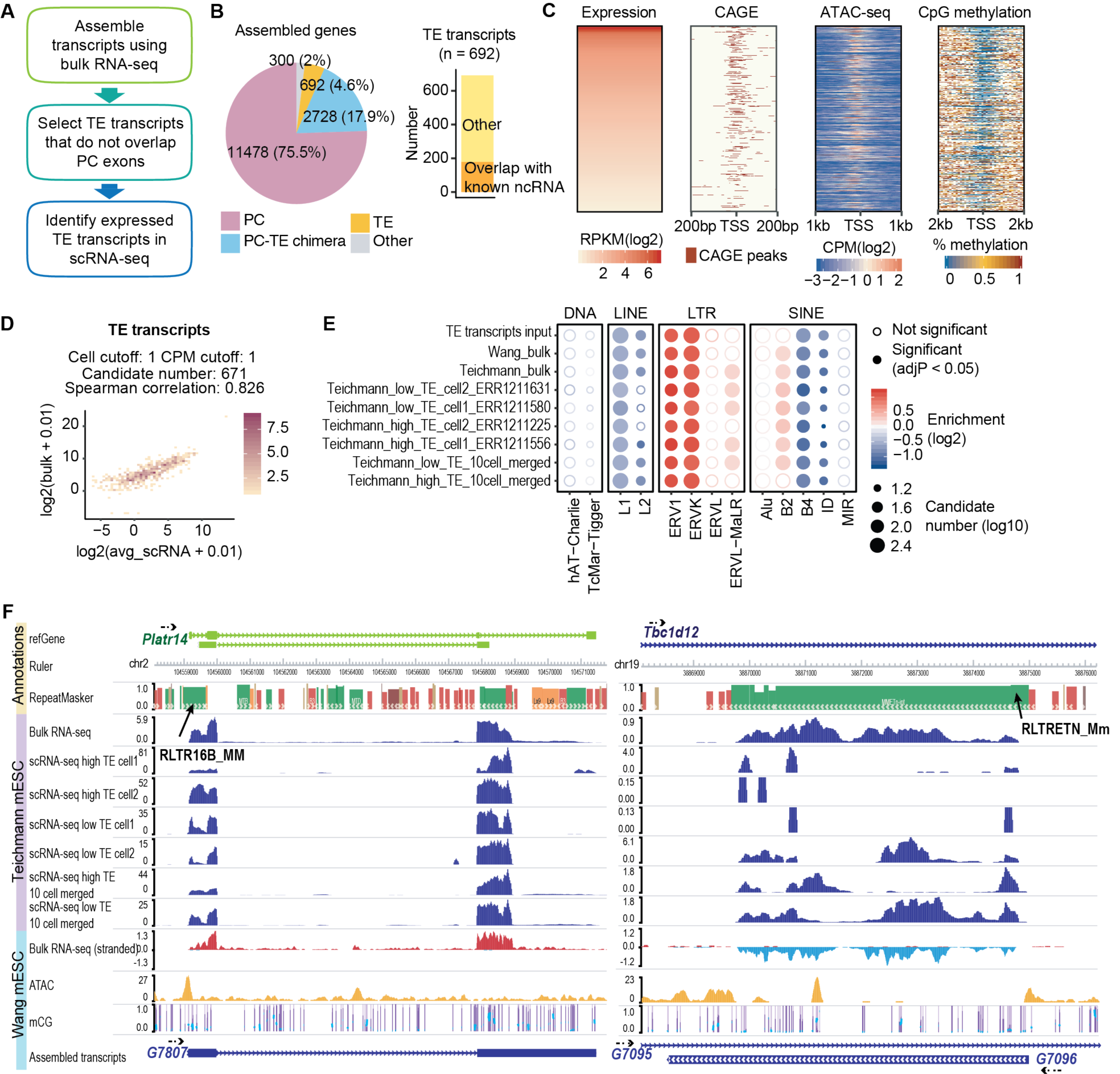
Transcript assembly improves scRNA-seq TE expression analysis. (A) Flowchart of scRNA-seq TE quantification pipeline. In short, transcript assembly was performed with bulk RNA-seq data and transcripts that overlap with TEs but not protein-coding exons were utilized for expression quantification in scRNA-seq data. (B) Transcript assembly using three mESC bulk RNA-seq data (Wang lab) yielded 692 TE transcripts. Among these TE transcripts, 179 overlap with ncRNAs annotated by refSeq. (C) FANTOM5 CAGE peaks, ATAC-seq signals and CpG methylation signals at the promoter region of TE transcripts with RPKM >=1 (Reads Per Kilobase Million). (D) Correlation between mESC bulk RNA-seq and averaged Smart-seq (Teichmann lab) signals at TE transcripts. (E) TE-family enrichment analysis using expressed TE transcripts. Enrichment of ERV elements was observed with both bulk RNA-seq and Smart-seq samples. (F) Examples of TE transcript. Assembled TE transcripts, uniquely mapped reads of mESC bulk RNA-seq, Smart-seq, merged Smart-seq, ATAC-seq and CpG methylation were included. Left: a TE transcript that initiates from RLTR16b_MM. This TE transcript overlaps Platr14, a long ncRNA known to impact the mESC differentiation-associated genes. Right: a TE transcript that initiates from RLTRETN_Mm. This transcript is largely composed of TEs and reflect the transcription unit of ERV.

Utilizing these newly generated transcript models, we recalculated the expression of TE transcripts and observed a much stronger correlation between mESC bulk and Smart-seq data (Fig. 2D). More importantly, we obtained much more consistent TE-family enrichment results between bulk and scRNA-seq and were able to identify the expression of ERV elements at single cells (Fig. 2E, F).

While Smart-seq based protocols generate deeper sequencing depth with reads covering the entire gene-body, other popular scRNA-seq strategies such as 10x Genomics Chromium, Drop-seq and SCRB-seq produce shallow sequencing with reads biased towards the 5’ or 3’ end of the RNA (Ramsköld et al. 2012; Soumillon et al. 2014; Macosko et al. 2015; Zheng et al. 2017). Counting reads at individual TEs using data with 5’ or 3’ signal enrichment will only capture TEs that are located at either end of the transcripts, thus biasing our understanding about TE expression. We reason that our approach can help to overcome this limitation by utilizing the annotation of full-length TE transcripts.

To support our reasoning, we analyzed the dataset from a previously published mESC differentiation study, in which single cell Smart-seq2, single cell SCRB-seq, and bulk RNA processed with SCRB-seq protocol were performed (Semrau et al. 2017). Using individual TEs as reference, we observed a severe discordance of TE expression between SCRB-seq and Smart-seq2, likely due to differences in signal distribution along the transcripts (Supplemental Fig. S5A). Conversely, using full-length TE transcripts as reference led to significantly improved signal correlations (Supplemental Fig. S5B). Moreover, quantifying TE expression at transcript level allowed us to recover the enrichment of ERVs from all datasets, whereas only Smart-seq2 showed ERV enrichments when counting at individual TEs (Supplemental Fig. S5C).

Taken together, these results suggest that our analysis approach is applicable not only to scRNA-seq data with high number of reads covering the entire transcript body, but also to other popular scRNA-seq strategies that feature shallow sequencing depth at the 3’ end of the transcripts.

### Dynamic TE expression in pre-implementation embryos

Encouraged by the results from the mESC data, we decided to apply our strategy to a more complex biological system: mouse embryogenesis. The dynamic regulation of epigenome during development not only fine-tunes protein-coding genes, but also allows specific TE expression at different developmental stages (Rowe and Trono 2011; Gifford et al. 2013; Gerdes et al. 2016; Rodriguez-Terrones and Torres-Padilla 2018; Deniz et al. 2019). Several recent studies utilized scRNA-seq to profile the transcription landscape of mouse embryos from zygote to early organogenesis, providing valuable resources for dissecting the dynamic expression of TEs.

To facilitate TE expression quantification, we performed transcript assembly using 37 bulk RNA-seq samples (Supplemental Table S1) that cover a range of tissues and developmental stages and obtained 5299 TE transcripts (Fig. 3A, B, Supplemental Fig. S6A, Supplemental Table S2). 770 of these transcripts overlap with known ncRNAs annotated by refSeq (Fig. 3A). Compared with assembled protein-coding transcripts, which show similar length and exon number as refSeq protein-coding gene annotations, assembled TE transcripts are shorter in length and possess fewer exons, a pattern consistent with annotated ncRNAs (Supplemental Fig. S6B, C).

**Figure 3.**
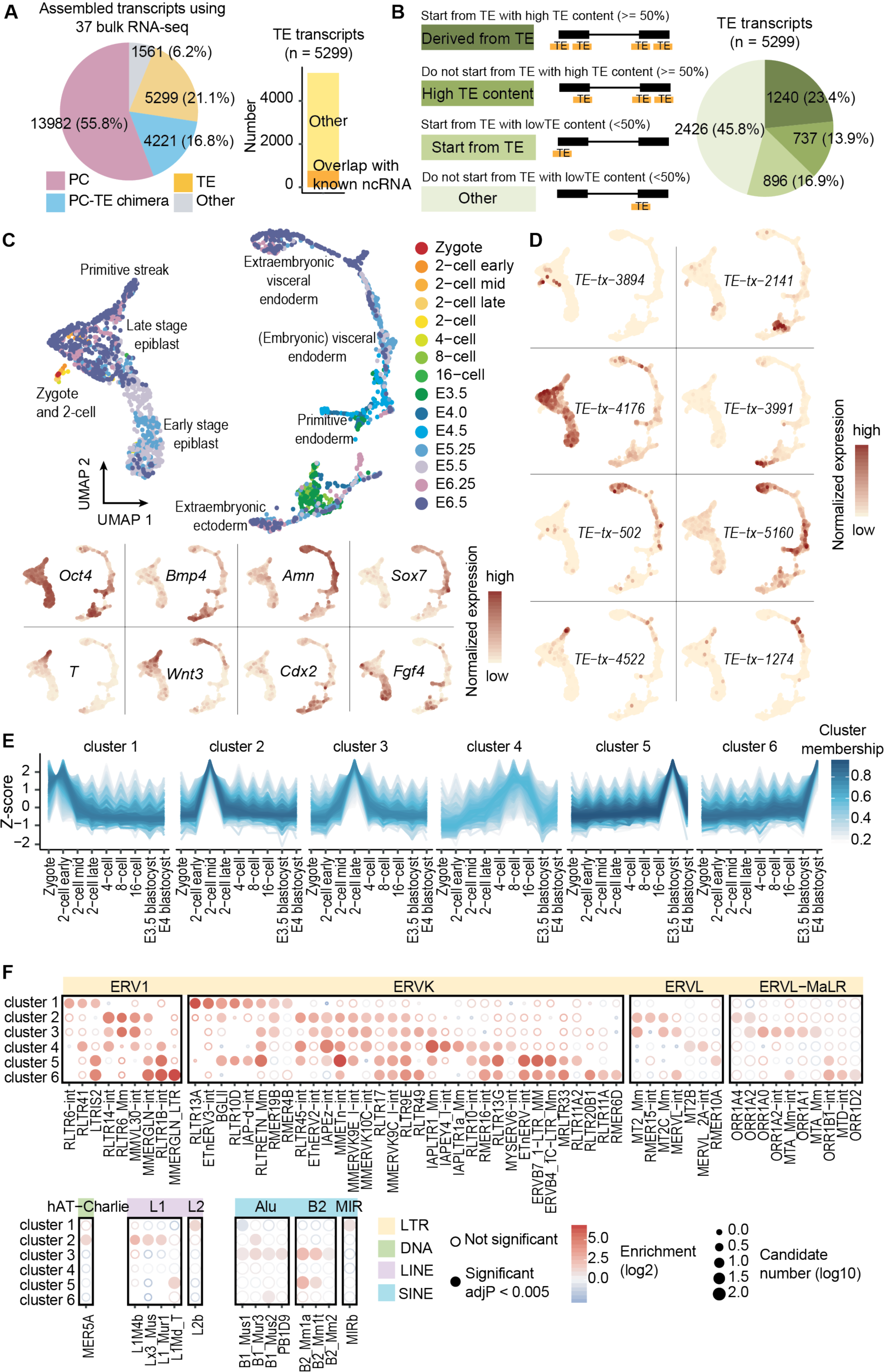
Dynamic TE expression in mouse pre-implantation embryos. (A) 5299 TE transcripts were constructed using 37 bulk RNA-seq samples. 770 of these TE transcripts overlap with ncRNAs annotated by refSeq. (B) Over half of all the assembled TE transcripts either initiate from TEs or have more than 50% of their exons composed of TEs. (C) Upper panel: UMAP of scRNA-seq data from mouse zygote to E6.5 embryos. Cells were colored based on developmental stages. Lower panel: expression of cell type specific markers. (D) Examples of developmental stage- and tissue-specific TE transcripts. (E) TE transcripts were grouped into 6 clusters based on their expression pattern across pre-implantation stages. (F) TE-subfamily enrichment analysis using TE transcripts within each of the 6 clusters.

Next, we analyzed three publicly available datasets where the transcription landscape of mouse embryos from zygote to gastrulation was profiled using Smart-seq derived protocols (Deng et al. 2014; Mohammed et al. 2017; Cheng et al. 2019). In these datasets, a significant number of reads overlap with TEs and about 3% of the total reads are mapped to TE transcripts (Supplemental Fig. S7A, B). After data integration and dimension reduction using the top 4000 variable features, we observed clear clustering patterns that were driven by cell type and developmental stage (Fig. 3C). Among the top 4000 variable features, 377 are TE transcripts, suggesting that the expression of TE transcripts could be cell-type- or developmental-stage-specific (Supplemental Fig. S7C-E). Indeed, we were able to observe TE transcript expression with strong tissue or stage specificity (Fig. 3D).

To further investigate the dynamics of TE transcription, we focused on pre-implantation stages, where high TE expression was documented. Due to the limited number of cells, scRNA-seq signals of each TE transcript across all the cells with the same developmental stage were averaged to reduce noise. Grouping TE transcripts based on their expression patterns across pre-implantation stages resulted in the following 6 clusters (Fig. 3E): TE transcripts that are maternally deposited (cluster1), TE transcripts that are up-regulated during minor and major waves of zygotic genome activation (clusters 2 and 3), TE transcripts that are upregulated during zygotic genome activation and keep accumulating till the blastocyst stage (cluster 4), TE transcripts that are up-regulated in the early- and mid-blastocyst stage (clusters 5 and 6).

We next performed TE enrichment analysis and observed that TE transcripts with distinct expression profiles tend to be enriched for different TE subfamilies (Fig. 3F). For instance, IAP elements are highly enriched in cluster 4, consistent with previous report that IAP expression initiates from the 2-cell stage, accumulates and then disappears at the blastocyst stage (Pikó et al. 1984; Poznanski and Calarco 1991; Svoboda et al. 2004). We also observed the enrichment of ERVL and ERVL-MaLR members in clusters 2 and 3, matching previous studies suggesting that ERVL and ERVL-MaLR members are highly expressed during 2-cell stage, constituting about 5% of the total transcripts (Kigami et al. 2003; Peaston et al. 2004; Svoboda et al. 2004). Furthermore, consistent with functional studies that L1 contributes to the entry and exit of 2-cell stage (Jachowicz et al. 2017; Percharde et al. 2018), we observed that the expression of L1 subfamily members peaks at the 2 cell stage. Moreover, transcription factor binding site analysis using a 500 bp window upstream of TE transcripts identified footprints of transcription factors that are crucial for mouse early embryogenesis such as Krupple-like factors, GABPA and ELF3 (Ristevski et al. 2004; Kageyama et al. 2006; Bialkowska et al. 2017), suggesting shared regulatory networks between TE transcripts and protein-coding genes (Supplemental Fig. S8).

### Tissue specific TE expression during mouse gastrulation and early organogenesis

Comparing to pre-implantation stages, TE expression during gastrulation and organogenesis is much less well studied and a comprehensive catalog of tissue specific TE transcripts is lacking. To address this, we analyzed a 10x scRNA-seq dataset where more than 100k cells were assayed using mouse E6.5 to E8.5 embryos (Pijuan-Sala et al. 2019). Comparing with mESC or mouse pre-implantation data analyzed in previous sections, this E6.5 to E8.5 10x data contains considerably less TE overlapping reads, with around 1% of the UMI mapping to TE transcripts (Supplemental Fig. S9A, B). Even though TE transcripts are in general lowly expressed and lack the extreme standardized variance observed at some protein-coding genes, they still constitute a small proportion of the top 1000 variable features that can be used to recapitulate the clustering pattern in the original study (Fig 4A, B, Supplemental Fig. S9C). More importantly, we were able to observe TE expression patterns that are enriched in small clusters of cells, suggesting that TE transcripts display considerable tissue specificity during these stages (Fig. 4C).

**Figure 4.**
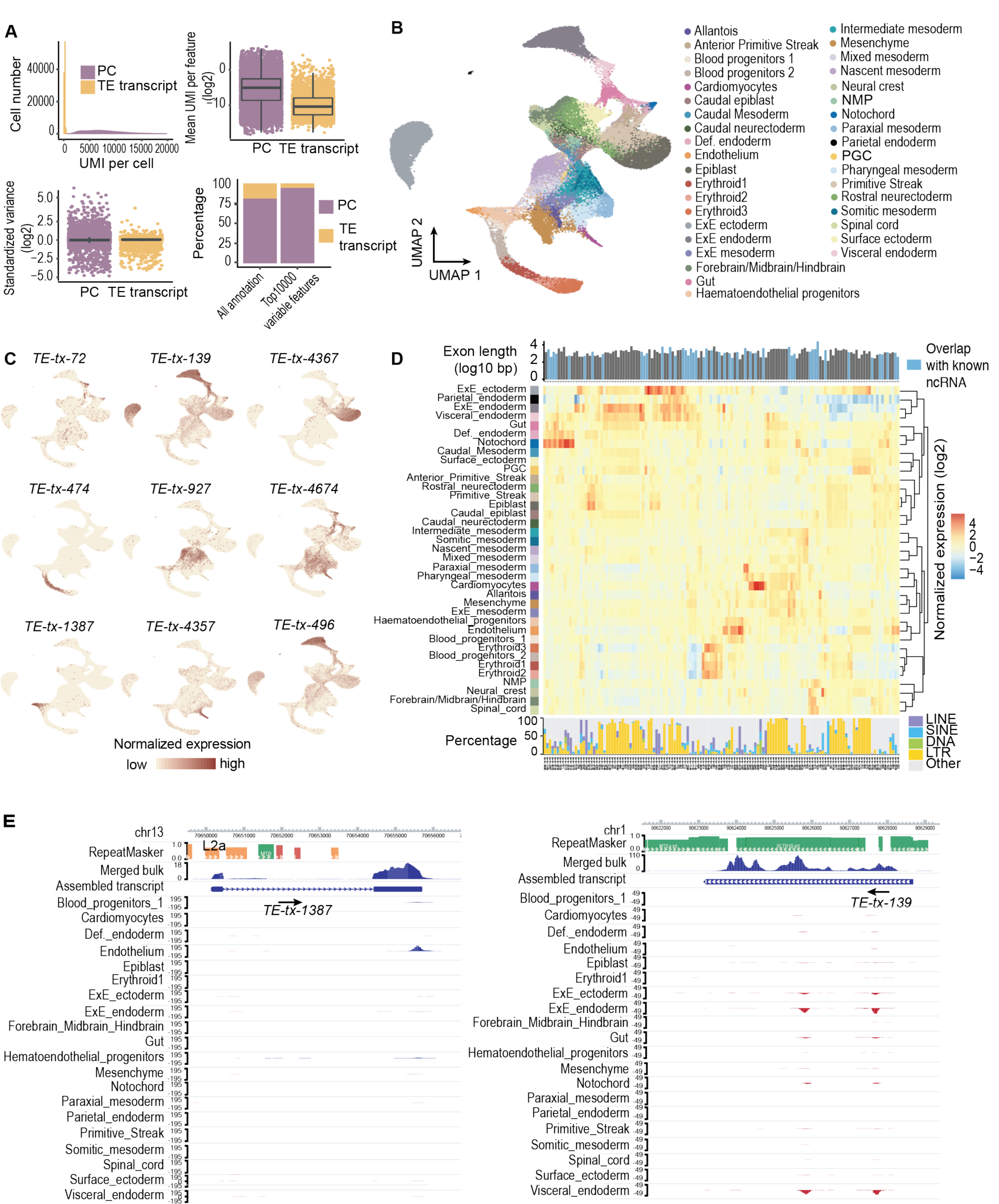
Tissue-specific TE expression during mouse gastrulation and early organogenesis. (A) Upper left: Fewer unique molecular identifiers (UMIs) were mapped to TE transcripts than to protein-coding genes. Upper right: the averaged expression level of TE transcripts across all the cells was lower compared to protein-coding genes. Lower left: TE transcripts lack the extreme standardized variance observed at protein-coding genes. Lower right: TE transcripts account for 55 of the top 1000 variable features. (B) UMAP of scRNA-seq data, cells were colored based on tissue information provided by the original study. (C) Examples of tissue specific TE transcripts. (D) Normalized expression pattern (center, heatmap) of 137 TE transcripts (columns) across 37 tissues (rows). Transcript length, annotation status (top, bar plot) and TE composition (bottom, bar plot) were shown for each TE transcript. (E) Genome browser view of two TE transcripts with strong tissue enrichment. Assembled TE transcripts, uniquely mapped reads of merged bulk RNA-seq (from 37 samples that were used for transcript assembly) and scRNA-seq signal for selected tissues were shown.

To systematically examine the dynamic TE expression, we obtained 137 TE transcripts with substantial tissue enrichment. Hierarchical clustering analysis using the expression of these TE transcripts showed that tissues with similar origins are grouped together (Fig. 4D). For instance, tissues within the hemato-endothelial lineage including hematoendothelial progenitors, endothelium, blood progenitors and erythroids are adjacent to each other, and tissues linked to the neuronal lineage including neuromesodermal progenitor, spinal cord, forebrain/midbrain/hindbrain and neural crest are clustered together.

Even though 10x reads were enriched at the 3’ end of the transcript, all TEs located along the transcripts were captured using our assembled full-length TE transcripts (Fig. 4E). Among the 137 TE transcripts, 82 initiate from TE or have more than 50% of their exonic sequences contributed by TEs (Fig. 4D, Supplemental Fig. S9D). Intriguingly, we observed that TE transcripts enriched in different tissues display distinct TE composition. For example, while TE transcripts enriched in hemato-lineage related tissues contain mostly non-TE sequences, TE transcripts enriched in extraembryonic ectoderm, extraembryonic endoderm, parietal endoderm and visceral endoderm are almost exclusively composed of LTRs. Furthermore, overlapping these TE transcripts with annotated ncRNAs revealed that 47 have been annotated by refSeq. Importantly, we observed that while these known ncRNAs tend to contain a lower percentage of TEs, transcripts that are mostly exclusively composed of TEs are largely un-annotated, demonstrating the value of our approach in capturing transcripts that originated from highly repetitive regions.

## Discussion

Current genome-wide TE expression quantification tools often count signal at individual TEs or TE subfamilies/families (Day et al. 2010; Jin et al. 2015; Lerat et al. 2017; Jeong et al. 2018; Yang et al. 2019; He et al. 2020; Jonsson et al. 2020). This strategy has been widely adopted in bulk RNA-seq and inspired similar analyses with scRNA-seq data (Göke et al. 2015; Ge 2017; Boroviak et al. 2018; Brocks et al. 2018; Yandim and Karakülah 2019). However, we caution that compared with bulk RNA-seq, a higher percentage of scRNA-seq reads are mapped to TEs, making it challenging to identify *bona fide* TE expression. Moreover, quantifying signal at individual TEs or TE subfamilies/families leads to the false impression that transcripts originated from repetitive regions are mostly composed of a single TE or TEs from the same subfamilies/families. While this is true for some well-studied examples such as full length ERVs, in most other cases, TEs only contribute to fragments of the full-length transcript and TEs from different families can be incorporated into the same transcript.

A major difference between the expression quantification of protein-coding genes and TEs is that the transcript structures of protein-coding genes are usually well annotated and readily available. Gene annotation guides expression analysis towards genomic regions with true signal and facilitates accurate expression quantification with scRNA-seq data. In this study, we demonstrated that transcripts constructed from bulk RNA-seq can serve as references for TE-containing ncRNAs and improve the accuracy of TE expression analysis in scRNA-seq data generated across multiple sequencing platforms. In comparison to individual TEs or TE subfamilies/families, TE transcripts more accurately reflect the natural transcription units. These transcripts contain not only previously annotated TE transcription units, but also novel ncRNAs that are partially composed of TEs. Out of the 5299 TE transcripts that we assembled, 98 closely resemble the well-studied transcription units of ERVs. These transcripts have more than 80% of their exonic sequences contributed by TEs. They start from 5’ LTR, transcribe through internal sequences and end at 3’ LTR. Notably, we also obtained another 104 TE transcripts that initiate from TEs and are within 100 bp away from FANTOM5 CAGE peaks (FANTOM Consortium and the RIKEN PMI and CLST (DGT) et al. 2014). 84 of these transcripts were not previously annotated, highlighting the value of our approach in identifying TEs that potentially function as promoters.

Using our analytical pipeline, we dissected the expression dynamics of TE transcripts during mouse early embryogenesis and identified 137 TE containing ncRNAs with strong tissue specificity. A close examination of these candidates revealed intricate interaction between TE transcripts and protein-coding genes. For instance, we were able to identify a ChIP-seq peak of Regulatory Factor X3 (RFX3) at the promoter region of the TE transcript *TE-tx-3856* (Supplemental Fig. 10A). RFX3 is a transcription factor essential for brain development (Baas et al. 2006; Benadiba et al. 2012; Magnani et al. 2015). Our observation that *TE-tx-3856* is highly expressed in mouse neuronal tissues suggests that this TE containing ncRNA is a potential downstream target of RFX3. In another interesting example, the TE transcript *TE-tx-3715* overlaps with *Sonic Hedgehog* (*Shh*), a secreted signaling molecule produced by the notochord (Placzek 1995; McMahon et al. 2003). Intriguingly, the expression pattern of *TE-tx-3715* strongly resembles that of *Shh*, indicating that they are under the control of a common regulatory circuit (Supplemental Fig. S10B). In addition, we also captured TE transcripts *TE-tx-3178* and *TE-tx-2841*, both of which are strongly expressed in the epiblast (Supplemental Fig. S10C). Both candidates initiate from TEs and were previously annotated as pluripotent associated transcripts (*Platr10* and *Platr14*) (Bergmann et al. 2015). Earlier reports suggested that *Platr10* transcript physically interacts with the promoter of pluripotent transcription factor *Sox2* while the depletion of *Platr14* alters the expression of differentiation- and development-associated genes in stem cells (Bergmann et al. 2015; Zhang et al. 2019). Our observation that *Platr10* and *Platr14* are expressed in the epiblast suggests that they may play similar roles during mouse early embryogenesis. Taken together, we dissected the dynamic TE expression during mouse early development and provided a curated list of promising TE candidates for future functional studies.

In summary, we established an effective TE quantification pipeline for scRNA-seq data and illustrated the dynamic TE expression during mouse early embryogenesis. In contrast to commonly used bulk RNA-seq tools that evaluate reads at single TEs or TE subfamilies/families, our pipeline emphasizes the importance of full-length TE transcript structures in scRNA-seq TE quantification. Furthermore, our work provides an initial set of TE transcript references during mouse early development through transcript assembly, laying the foundation for future work on constructing a more comprehensive TE transcript database across tissues and developmental stages. Additionally, exploring alternative quantification methods for ambiguously mapped reads, such as using expectation–maximization algorithm or Bayesian-based framework, as well as developing isoform-specific quantification tools for TE-protein-coding-gene chimeras will further expand the TE analysis toolkit for scRNA-seq and greatly advance our knowledge on the expression and the function of TE transcripts.

## Methods

### Data processing and signal quantification of bulk RNA-seq datasets

Raw sequencing files were downloaded from NCBI Sequence Read Archive and EMBL-EBI ArrayExpress (Supplemental Table S1) and aligned to the mouse (mm10) or human (hg38) genomes using STAR. To retain reads derived from repetitive regions, “--outFilterMultimapNmax” was set to 500. To facilitate downstream transcript assembly “--outSAMattributes” was set to “NH HI NM MD XS AS”. After alignment, signal quantification at regions of interests was performed using featureCount. Reads aligned to multiple locations or overlapping multiple features were evenly distributed by enabling “-O -M --fraction”.

### Data processing and signal quantification of scRNA-seq datasets

scRNA-seq data generated with Smart-seq derived protocols were processed and quantified using the same procedures as bulk RNA-seq data. scRNA-seq data generated with the other protocols were processed using zUMIs with the following modifications: 1) To retain reads derived from repetitive regions, “--outFilterMultimapNmax” was set to 500 during STAR alignment. 2) To quantify reads that were mapped to multiple locations or features, “allowMultiOverlap”and “countMultiMappingReads” were set to TRUE for function “.runFeatureCount”. Bam files with cell barcode, UMI and the name of overlapping features were reported. 3) A customized R script was used to process the bam file generated in step2. Reads that were mapped to multiple locations or features were evenly distributed. UMIs were then evaluated at each feature.

Reads of the scRNA-seq datasets from mouse zygote to gastrulation (Smart-seq derived protocols) were quantified at protein-coding genes (refSeq annotation, n= 20779) and TE transcripts (assembled from bulk RNA-seq, n=5299). Only cells that had 200 to 18000 features and less than 10% mitochondria reads were kept. To remove batch effect and visualize all three datasets in the same UMAP, data integration was performed using Seurat with the top 4000 variable features. Cell type was determined using the stage information provided by the original studies, the expression patterns of cell type specific markers and Seurat clustering results.

UMIs of the 10x scRNA-seq dataset from mouse gastrulation to early organogenesis were quantified at protein-coding genes (refSeq annotation, n= 20779) and TE transcripts (assembled from bulk RNA-seq, n=5299). Sample_25 was removed due to higher batch effect. Only cells that have more than 200 features and were annotated by the original study were kept. Cell type information provided by the original study was utilized for identifying tissue specific markers. The 137 TE transcripts with strong tissue enrichment were obtained by combining and filtering Seurat-defined markers and customized markers. Seurat-defined markers were obtained by running “FindAllMarkers” with “only.pos = T, min.pct = 0.15” and selecting for TE transcripts with adjusted p-value < 0.05. Customized markers were obtained by identifying TE transcripts with at least 1 UMI in at least 10% of the cells in any tissue and selecting candidates that were expressed in less than 4 tissues. After combining Seurat-defined markers and customized markers, manual curation was performed to remove candidates that were highly expressed in large number of tissues or with sub-optimal transcript structures.

### TE transcript clustering in mouse pre-implantation stages

Due to the limited number of cells, scRNA-seq signals of each TE transcript across all the cells with the same developmental stage were averaged to reduce noise. TE transcripts were then grouped into 6 clusters using soft clustering (R package TCseq) based on their expression patterns across pre-implantation stages.

### Constructing TE transcripts

Transcript assembly of each RNA-seq sample was performed using StringTie2. “-j 2 -s 5 -f 0.05 -c 2” was used to improve the specificity of the assembly results. To generate the master reference file, assembled transcripts from multiple RNA-seq samples were then merged using TACO with default parameters. Transcripts that overlapped with TEs but not the exons of refSeq protein-coding genes were named as TE transcripts and utilized for TE expression quantification.

### TE subfamily/family enrichment analysis

For each TE-family, its enrichment was calculated using the following equation: The observed frequency of TEs belonging to this family in all candidates divided by the expected frequency of TEs belonging to this family in genomic regions that do not overlap with protein-coding genes. The significance for the observed frequency was calculated with Fisher’s exact test and corrected for multiple testing with the Benjamini–Hochberg method. Only TE families with more than 20 members in the candidates and more than 100 members in the background were included in the figures. TE-subfamily enrichment analysis was performed similarly. Only TE-subfamilies that were significantly enriched in the candidates, had more than 10 members in the candidates and more than 100 members in the background and were plotted.

### Publicly available datasets utilized in this study

Descriptions and accession IDs of all the datasets used in this manuscript are provided in Supplemental Table S1.

### Data Access

All of the datasets analyzed in the paper are currently available in public databases, accession IDs and descriptions are listed in the Supplemental Table S1.

## Acknowledgements

We thank Kara Quaid and Shuonan He for comments on the manuscript, and all the Ting Wang lab members for discussions and critical inputs.

