## Supplementary materials for "Transcript assembly improves expression quantification of transposable elements in single cell RNA-seq data"

Shao and Wang, 2020

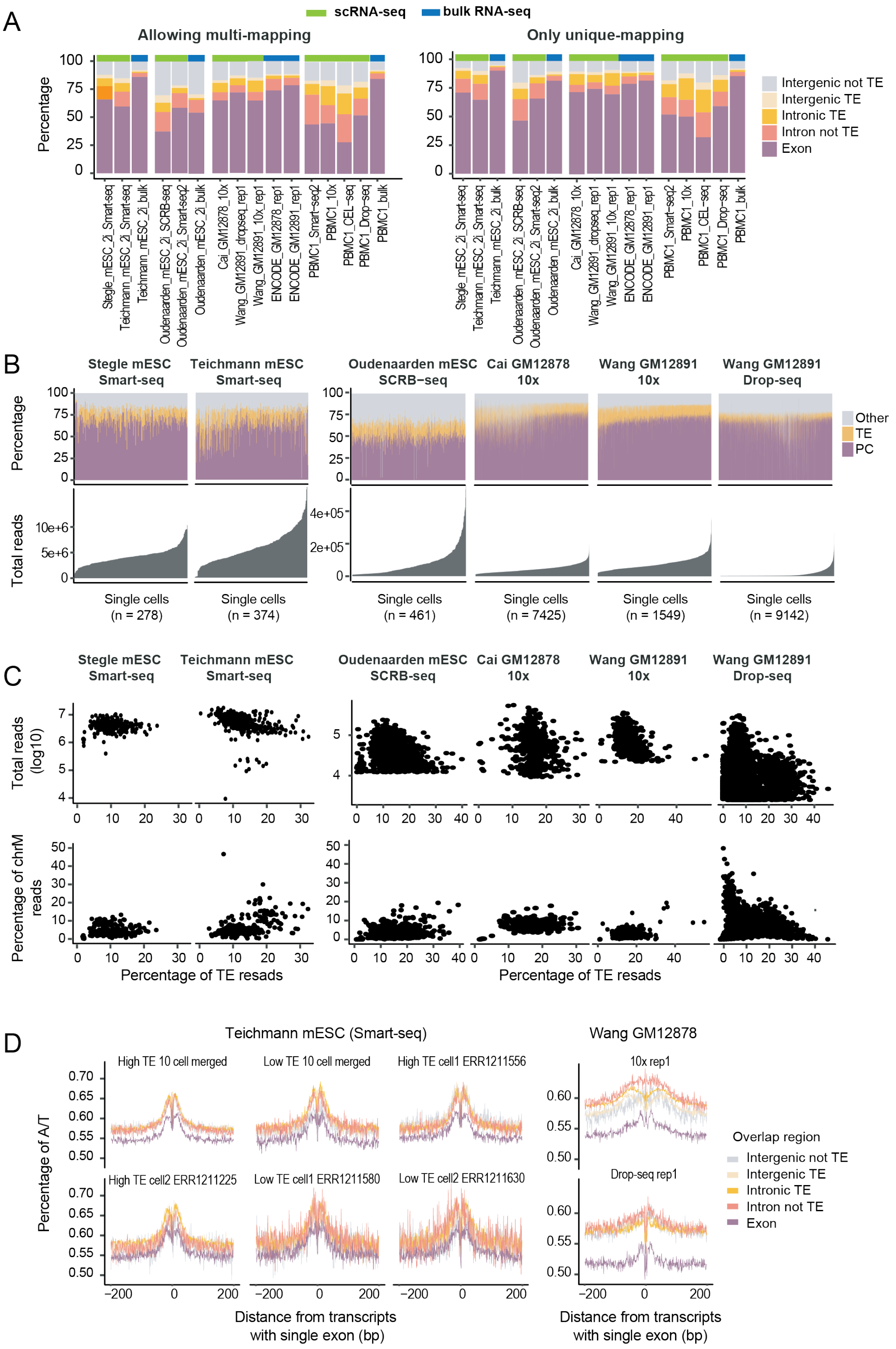
A
scRNA-seq
bulk RNA-seq
Allowing multi-mapping
Only unique-mapping
Percentage
Intergenic not TE
Intergenic TE
Intronic TE
Intron not TE
Exon
Stegle_mESC_2i_Smart-seq
Teichmann_mESC_2i_Smart-seq
Teichmann_mESC_2i_bulk
Oudenaarden_mESC_2i_SCRB-seq
Oudenaarden_mESC_2i_Smart-seq2
Oudenaarden_mESC_2i_bulk
Cai_GM12878_10x
Wang_GM12891_dropseq_rep1
Wang_GM12891_10x_rep1
ENCODE_GM12878_rep1
ENCODE_GM12891_rep1
PBMC1_Smart-seq2
PBMC1_10x
PBMC1_CEL-seq
PBMC1_Drop-seq
PBMC1_bulk
B
Stegle mESC Smart-seq
Teichmann mESC Smart-seq
Oudenaarden mESC SCRB−seq
Cai GM12878 10x
Wang GM12891 10x
Wang GM12891 Drop-seq
Other
TE
PC
Total reads
Single cells (n = 278)
Single cells (n = 374)
Single cells (n = 461)
Single cells (n = 7425)
Single cells (n = 1549)
Single cells (n = 9142)
C
Stegle mESC Smart-seq
Teichmann mESC Smart-seq
Oudenaarden mESC SCRB-seq
Cai GM12878 10x
Wang GM12891 10x
Wang GM12891 Drop-seq
Total reads (log10)
Percentage of chrM reads
Percentage of TE resads
D
Teichmann mESC (Smart-seq)
Wang GM12878
High TE 10 cell merged
Low TE 10 cell merged
High TE cell1 ERR1211556
High TE cell2 ERR1211225
Low TE cell1 ERR1211580
Low TE cell2 ERR1211630
10x rep1
Drop-seq rep1
Percentage of A/T
Overlap region
Intergenic not TE
Intergenic TE
Intronic TE
Intron not TE
Exon
Distance from transcripts with single exon (bp)

**Supplemental Figure S1 Evaluation of scRNA-seq reads that were mapped to TEs**

(A) Distribution of bulk RNA-seq and scRNA-seq reads in five non-overlapping genomic regions: protein-coding exons, TEs within the introns of protein-coding genes, other intronic regions of protein-coding genes, intergenic TEs, and other intergenic regions. Left: Reads mapped to multiple locations were included during signal quantification. Right: Only uniquely mapped reads were utilized.

(B) Distribution of scRNA-seq reads in single cells. Cells were arranged based on sequencing depth. PC: protein-coding exons. TE: transposable elements that do not overlap with protein-coding exons. Other: other genomic locations.

(C) Top: correlation between the percentage of TE reads and the sequencing depth of single cells. Bottom: correlation between the percentage of TE reads and that of the mitochondria reads.

(D) Regions enriched for un-spliced scRNA-seq reads tend to be flanked by AT-rich sequences. Transcripts assembly was performed using single cell data and transcripts with single exon were separated into 5 groups based on their genomic locations. A/T percentage was calculated at sequences flanking these transcripts.

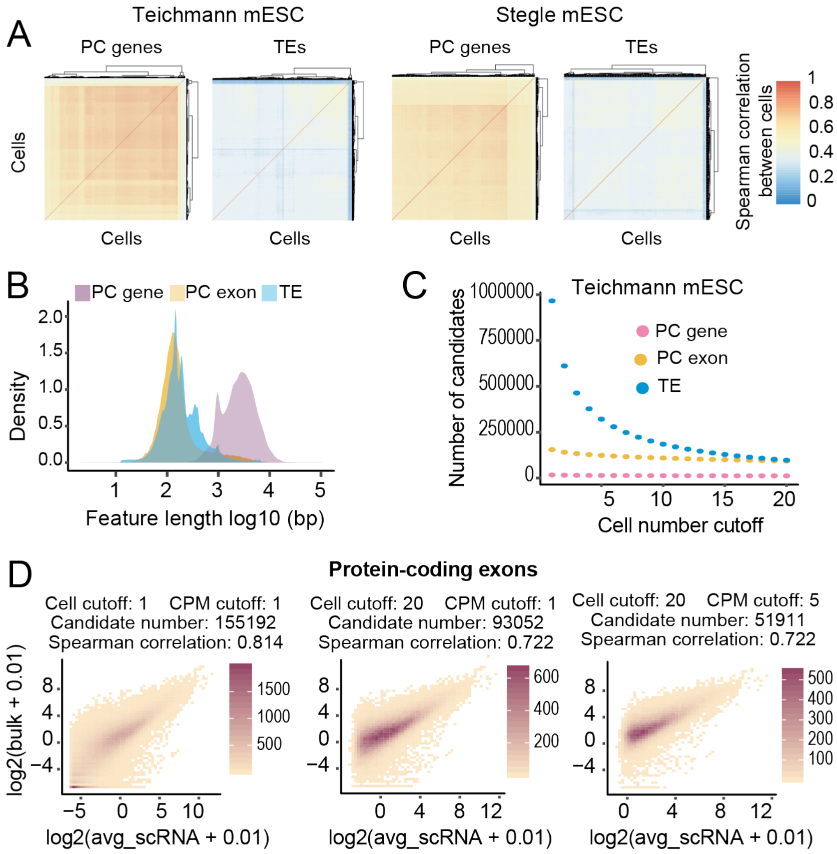

**Supplemental Figure S2 Length discrepancy between TEs and protein-coding genes cannot sufficiently explain the higher number of false positive candidates at TEs**

(A) Spearman correlation between cells using TEs or protein-coding genes as references. Higher correlation was observed at protein-coding genes.

(B) The length of full transcripts of protein-coding genes, exons of protein-coding genes and TEs.

(C) Number of candidates as a function of cell number cutoff (the minimum number of cells each candidate is expressed in. Expression cutoff: CMP >=1). Although majority of protein-coding genes and protein-coding exons were consistently detected, a large number of TE candidates were detected in less than 10 cells.

(D) Correlation between bulk RNA-seq and averaged Smart-seq signal at protein-coding exons (Teichmann lab, mESC). Cell cutoff: the minimum number of cells each candidate is expressed in. CPM cutoff: the minimum CPM value for one candidate to be considered as expressed.

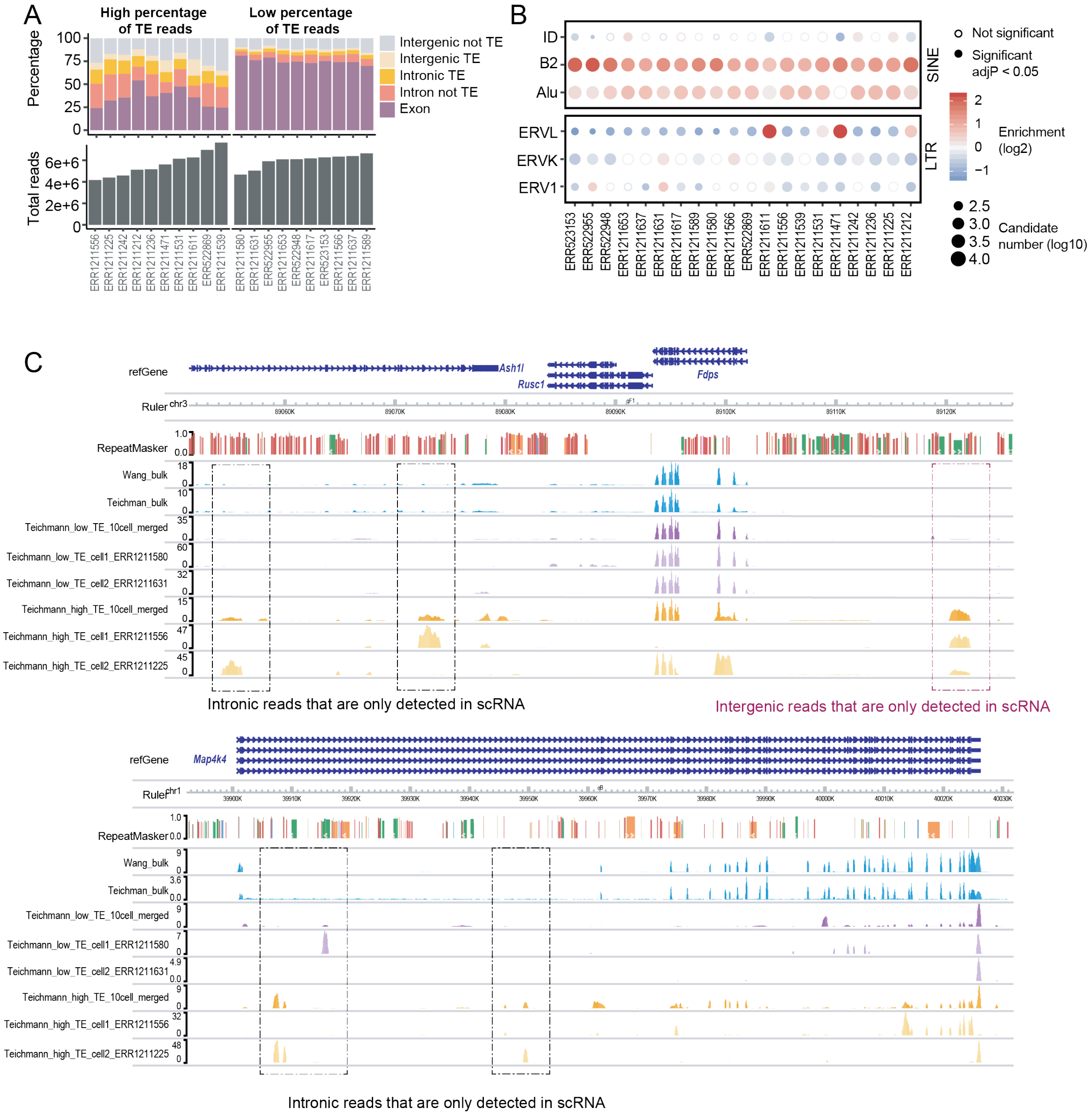
A
High percentage of TE reads
Low percentage of TE reads
Percentage
Total reads
Intergenic not TE
Intergenic TE
Intronic TE
Intron not TE
Exon
B
ID
B2
Alu
SINE
ERVL
ERVK
ERV1
LTR
Not significant
Significant adjP < 0.05
Enrichment (log2)
Candidate number (log10)
C
refGene
Ruler
RepeatMasker
Ash1l
Rusc1
Fdps
Map4k4
Wang_bulk
Teichman_bulk
Teichmann_low_TE_10cell_merged
Teichmann_low_TE_cell1_ERR1211580
Teichmann_low_TE_cell2_ERR1211631
Teichmann_high_TE_10cell_merged
Teichmann_high_TE_cell1_ERR1211556
Teichmann_high_TE_cell2_ERR1211225
Intronic reads that are only detected in scRNA
Intergenic reads that are only detected in scRNA
Intronic reads that are only detected in scRNA

**Supplemental Figure S3 Examples of single cells with different percentage of TE reads**

(A) Examples of single cells with similar sequencing depth but different percentage of TE reads (Teichmann lab, mESC). Top: distribution of scRNA-seq reads at 5 non-overlapping genomic regions in single cells. Bottom: sequencing depth per cell.

(B) TE-family enrichment analysis at single cells with different percentage of TE reads (Teichmann lab, mESC).

(C) Genome browser view of mESC bulk RNA-seq, Smart-seq of four single cells and merged Smart-seq. Only uniquely mapped reads are shown.

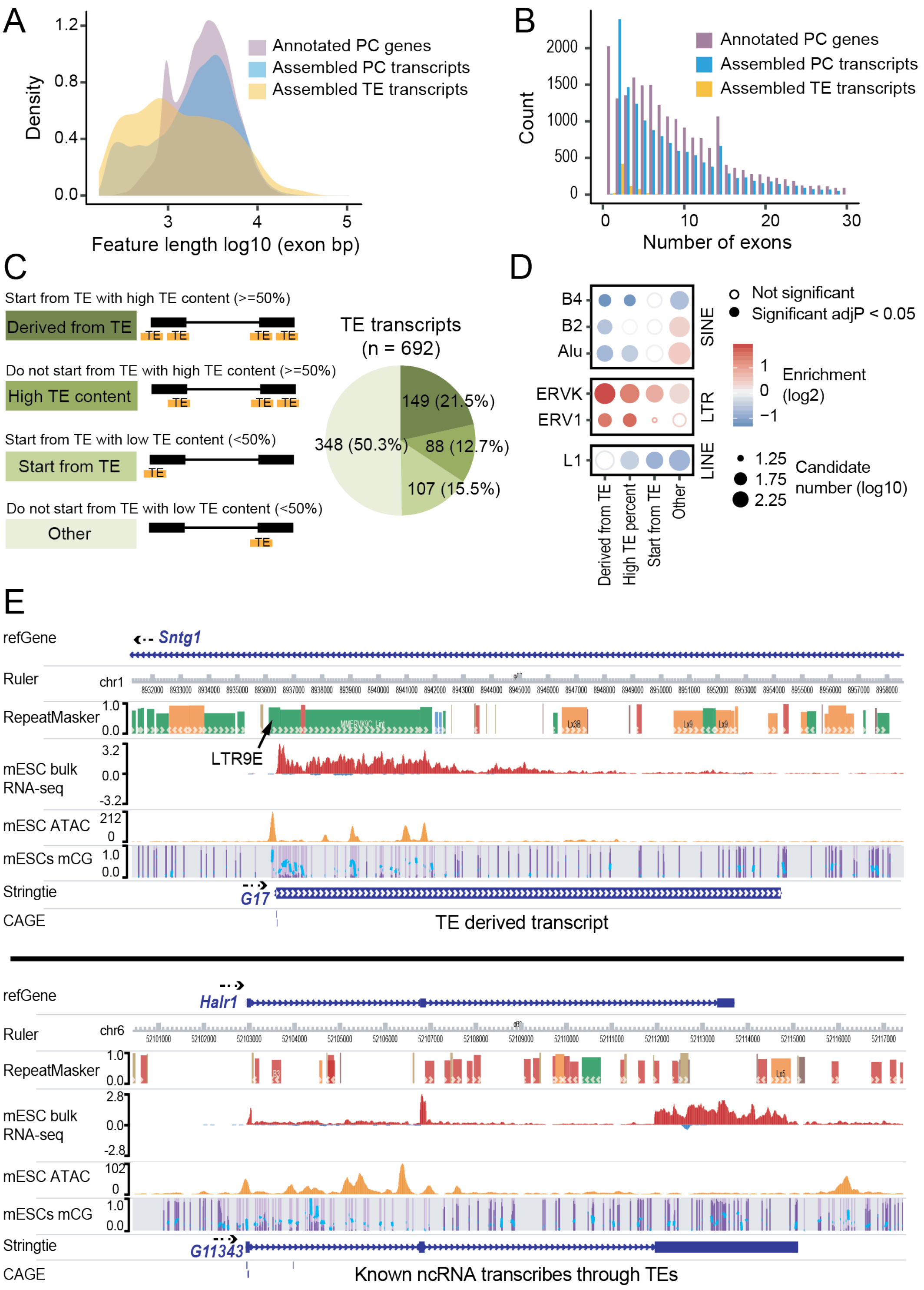
A
Density
Feature length log10 (exon bp)
Annotated PC genes
Assembled PC transcripts
Assembled TE transcripts
B
Count
Number of exons
Annotated PC genes
Assembled PC transcripts
Assembled TE transcripts
C
Start from TE with high TE content (>=50%)
Derived from TE
Do not start from TE with high TE content (>=50%)
High TE content
Start from TE with low TE content (<50%)
Start from TE
Do not start from TE with low TE content (<50%)
Other
TE transcripts
(n = 692)
149 (21.5%)
88 (12.7%)
107 (15.5%)
348 (50.3%)
D
B4
B2
Alu
ERVK
ERV1
L1
SINE
LTR
LINE
Derived from TE
High TE percent
Start from TE
Other
Not significant
Significant adjP < 0.05
Enrichment
(log2)
Candidate
number (log10)
E
refGene
Sntg1
Ruler
RepeatMasker
LTR9E
mESC bulk
RNA-seq
mESC ATAC
mESCs mCG
Stringtie
G17
CAGE
TE derived transcript
refGene
Halr1
Ruler
RepeatMasker
mESC bulk
RNA-seq
mESC ATAC
mESCs mCG
Stringtie
G11343
CAGE
Known ncRNA transcribes through TEs

**Supplemental Figure S4 Evaluation of TE transcripts assembled from mESC bulk RNA-seq data**

(A) The length of protein-coding genes annotated by refSeq, assembled transcripts that overlap with annotated protein-coding genes and assembled TE transcripts. Only exonic regions were considered.

(B) The number of exons of protein-coding genes annotated by refSeq, assembled transcripts that overlap with annotated protein-coding genes and assembled TE transcripts.

(C) Around half of all the assembled TE transcripts either initiate from TEs or have more than 50% of their exons composed of TEs.

(D) TE-family enrichment analysis at different classes of TE transcripts. TE transcripts that initiate from TEs or have more than 50% of their exons composed of TEs are enriched for LTR elements.

(E) Genome browser view of bulk RNA-seq, ATAC-seq, CpG methylation, assembled TE transcripts (Wang lab, mESC) and FANTOM5 CAGE peaks. Only uniquely mapped reads are shown.

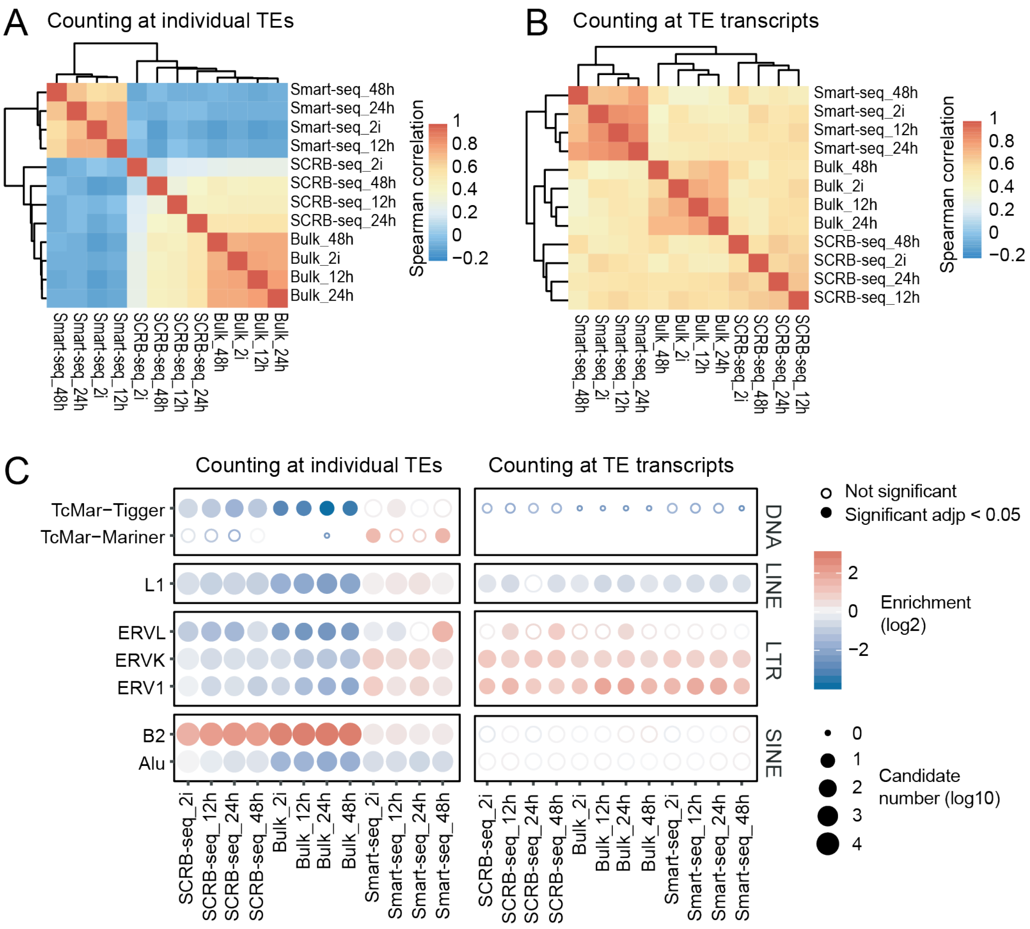

**Supplemental Figure S5 Quantifying signal at TE transcripts is applicable to scRNA-seq with 3' end signal enrichment**

(A) and (B) Spearman correlation of datasets from a previously published mESC differentiation study, in which single cell Smart-seq2, single cell SCRB-seq and bulk RNA processed with SCRB-seq protocol were performed. Using full-length TE transcripts as reference led to significantly improved signal correlation between SCRB-seq and Smart-seq2 samples.

(C) TE-family enrichment analysis. Quantifying TE expression at transcript level allowed us to recover the enrichment of ERVs from all datasets, whereas only Smart-seq2 showed ERV enrichments when counting at individual TE.

99

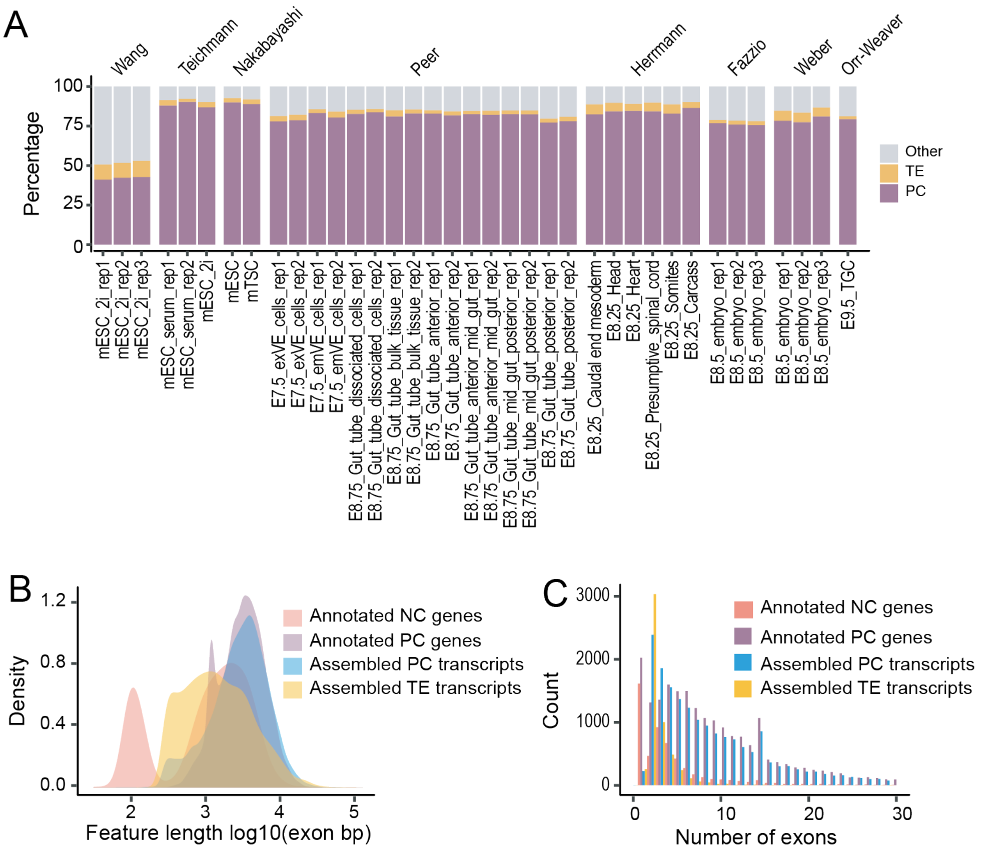

**Supplemental Figure S6 Evaluation of TE transcripts assembled from 37 bulk RNA-seq**

(A) Distribution of mappable reads in 37 bulk RNA-seq samples. PC: protein-coding exons defined by refSeq. TE: transposable elements that do not overlap with protein-coding exons. Other: other genomic locations.

(B) The length of protein-coding genes annotated by refSeq, noncoding genes annotated by refSeq, assembled transcripts that overlap with annotated protein-coding genes and assembled TE transcripts. Only exonic regions were considered.

(C) The number of exons of protein-coding genes annotated by refSeq, noncoding genes annotated by refSeq, assembled transcripts that overlap with annotated protein-coding genes and assembled TE transcripts.

112

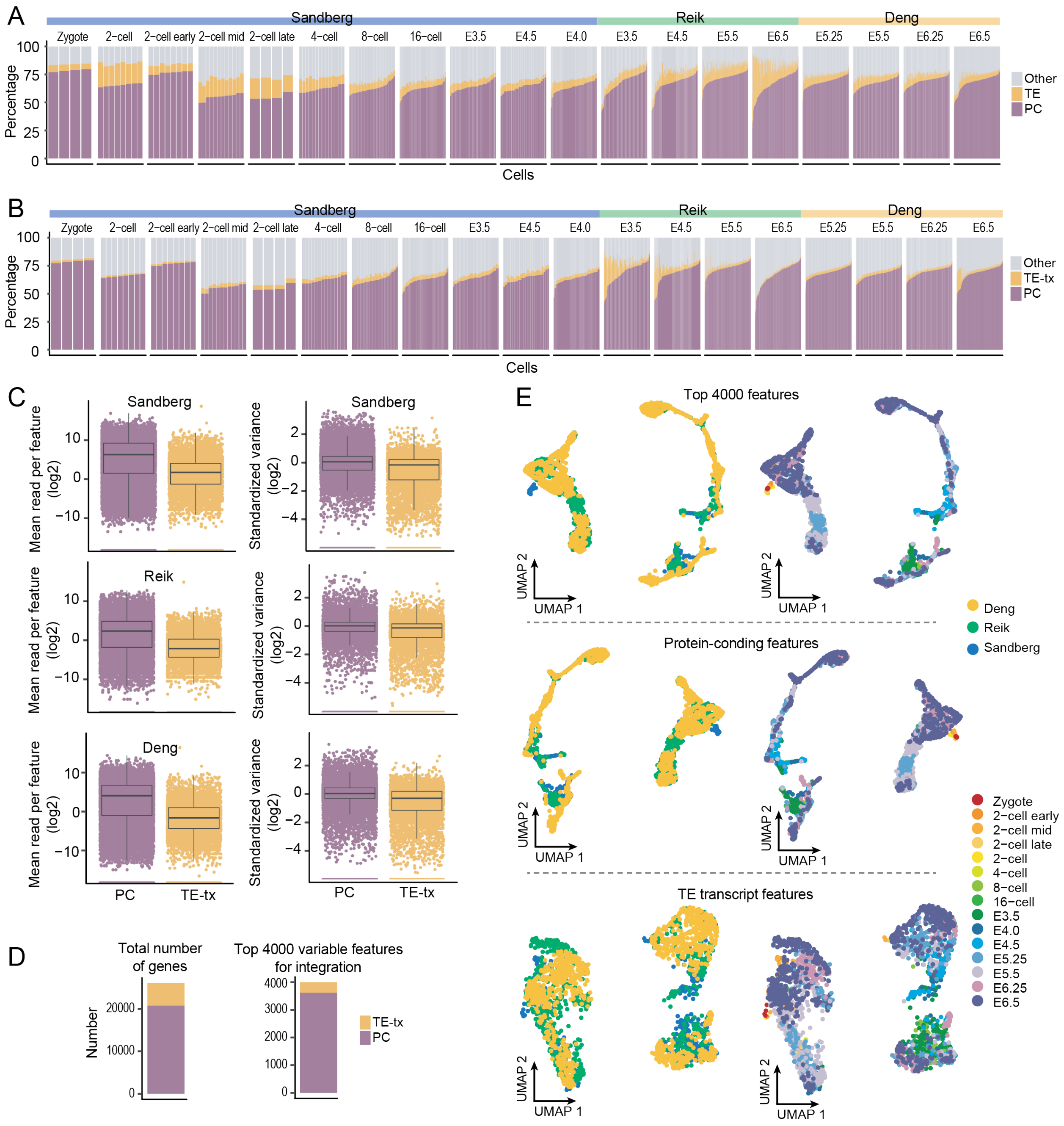

**Supplemental Figure S7 Evaluation of mouse scRNA-seq data from zygote to gastrulation**

(A) and (B) Distribution of mappable reads. PC: protein-coding exons defined by refSeq. TE: transposable elements that do not overlap with protein-coding exons. TE-tx: TE transcripts. Other: other genomic locations.

(C) Left: averaged signal of protein-coding genes and TE transcripts across all the cells. Right: standardized variance of protein-coding genes and TE transcripts.

(D) The number of TE transcripts and protein-coding genes in the top 4000 variable features. Left: 5299 TE transcripts and 20779 protein-coding genes were used for scRNA-seq signal quantification. Right: among the top 4000 variable features that were used for scRNA-seq dimension reduction and integration, 377 are TE transcripts.

(E) UMAP of scRNA-seq data. scRNA-seq dimension reduction and integration were performed using the top 4000 variable features (top), the top variable features that are protein-coding genes (middle, n = 3623), or the top variable features that are TE transcripts (bottom, n = 377).

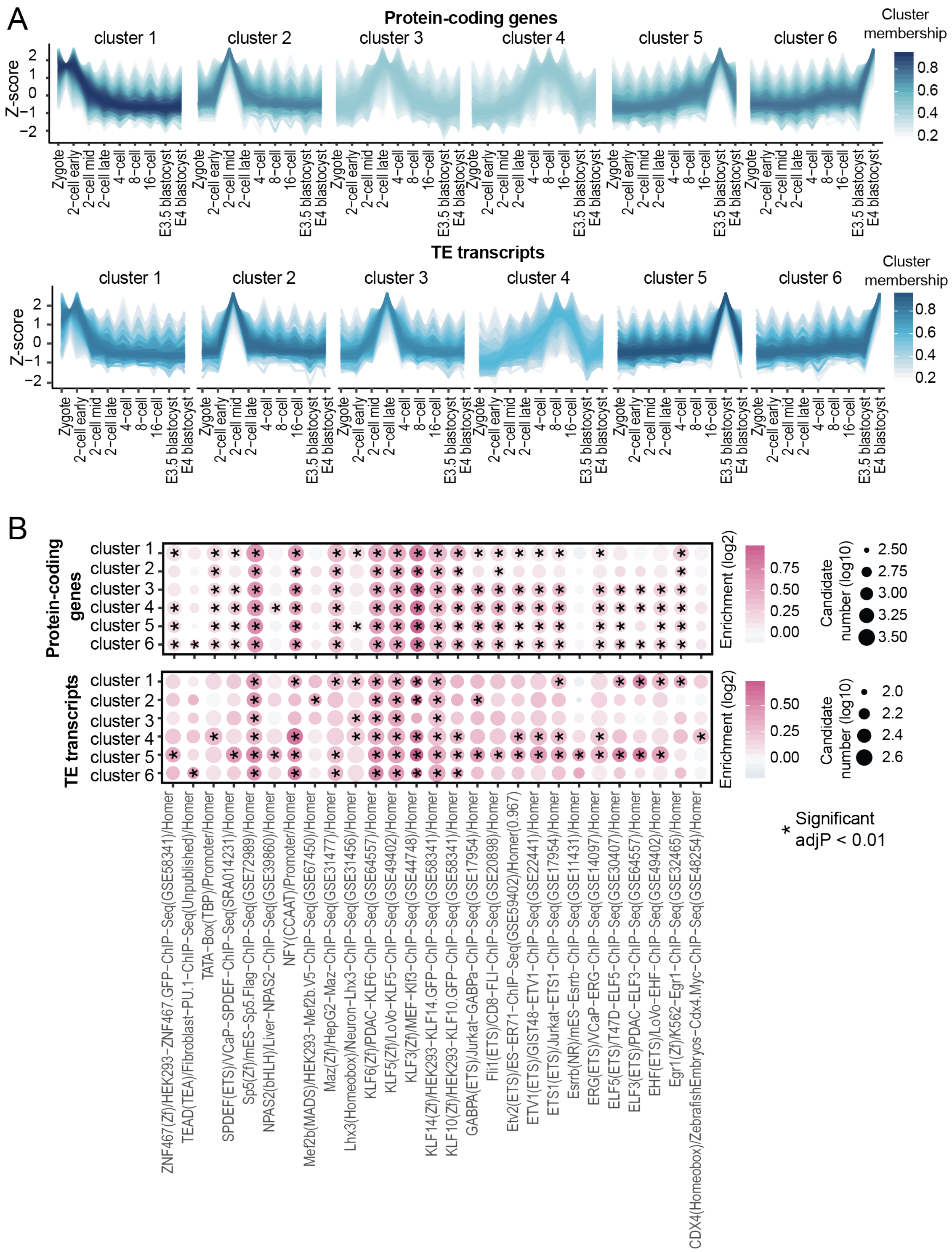
A
Protein-coding genes
cluster 1
cluster 2
cluster 3
cluster 4
cluster 5
cluster 6
Z-score
Cluster membership
Zygote
2−cell early
2−cell mid
2−cell late
4−cell
8−cell
16−cell
E3.5 blastocyst
E4 blastocyst
TE transcripts
B
Protein-coding genes
TE transcripts
Enrichment (log2)
Candidate number (log10)
* Significant adjP < 0.01
ZNF467(Zf)/HEK293−ZNF467.GFP−ChIP−Seq(GSE58341)/Homer
TEAD(TEA)/Fibroblast−PU.1−ChIP−Seq(Unpublished)/Homer
TATA−Box(TBP)/Promoter/Homer
SPDEF(ETS)/VCaP−SPDEF−ChIP−Seq(SRA014231)/Homer
Sp5(Zf)/mES−Sp5.Flag−ChIP−Seq(GSE72989)/Homer
NPAS2(bHLH)/Liver−NPAS2−ChIP−Seq(GSE39860)/Homer
NFY(CCAAT)/Promoter/Homer
Mef2b(MADS)/HEK293−Mef2b.V5−ChIP−Seq(GSE67450)/Homer
Maz(Zf)/HepG2−Maz−ChIP−Seq(GSE31477)/Homer
Lhx3(Homeobox)/Neuron−Lhx3−ChIP−Seq(GSE31456)/Homer
KLF6(Zf)/PDAC−KLF6−ChIP−Seq(GSE64557)/Homer
KLF5(Zf)/LoVo−KLF5−ChIP−Seq(GSE49402)/Homer
KLF3(Zf)/MEF−Klf3−ChIP−Seq(GSE44748)/Homer
KLF14(Zf)/HEK293−KLF14.GFP−ChIP−Seq(GSE58341)/Homer
KLF10(Zf)/HEK293−KLF10.GFP−ChIP−Seq(GSE58341)/Homer
GABPA(ETS)/Jurkat−GABPa−ChIP−Seq(GSE17954)/Homer
Fli1(ETS)/CD8−FLI−ChIP−Seq(GSE20898)/Homer
Etv2(ETS)/ES−ER71−ChIP−Seq(GSE59402)/Homer(0.967)
ETV1(ETS)/GIST48−ETV1−ChIP−Seq(GSE22441)/Homer
ETS1(ETS)/Jurkat−ETS1−ChIP−Seq(GSE17954)/Homer
Esrrb(NR)/mES−Esrrb−ChIP−Seq(GSE11431)/Homer
ERG(ETS)/VCaP−ERG−ChIP−Seq(GSE14097)/Homer
ELF5(ETS)/T47D−ELF5−ChIP−Seq(GSE30407)/Homer
ELF3(ETS)/PDAC−ELF3−ChIP−Seq(GSE64557)/Homer
EHF(ETS)/LoVo−EHF−ChIP−Seq(GSE49402)/Homer
Egr1(Zf)/K562−Egr1−ChIP−Seq(GSE32465)/Homer
CDX4(Homeobox)/ZebrafishEmbryos−Cdx4.Myc−ChIP−Seq(GSE48254)/Homer

**Supplemental Figure S8 Dynamic expression of TE transcripts and protein-coding genes during mouse pre-implantation stages**

(A) TE transcripts and protein-coding genes were grouped into 6 clusters based on their expression patterns across pre-implantation stages.

(B) Enrichment of known motifs at the promoter region of TE transcripts and protein-coding genes. HOMER2 was run with the promoter sequences of TE transcripts and protein-coding genes (500bp upstream of the transcription start site), known motifs that are shared between TE transcripts and protein-coding genes were shown.

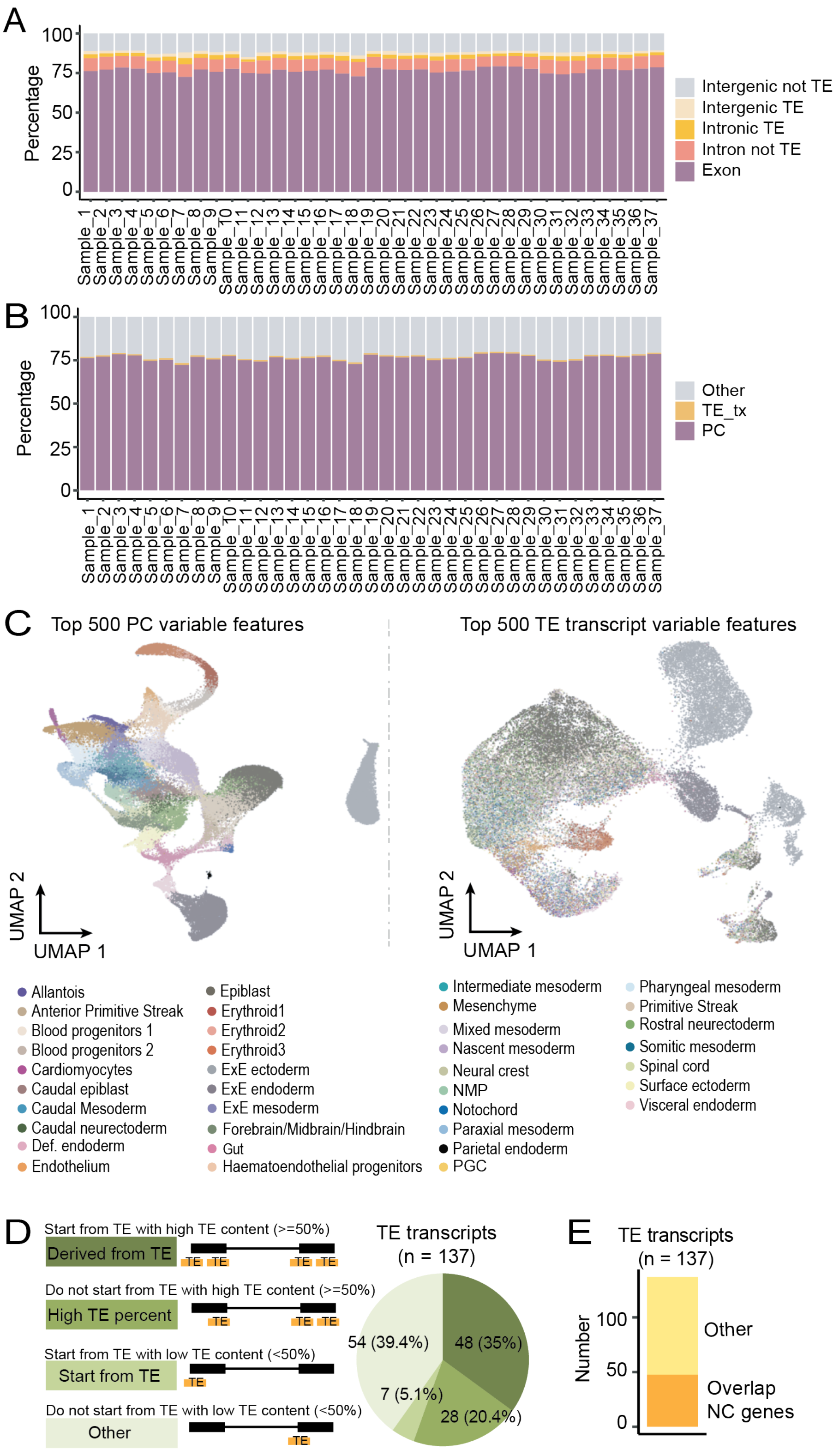
A
Percentage
100
75
50
25
0
Intergenic not TE
Intergenic TE
Intronic TE
Intron not TE
Exon
Sample_1 … Sample_37
B
Percentage
Other
TE_tx
PC
C
Top 500 PC variable features
Top 500 TE transcript variable features
UMAP 2
UMAP 1
Allantois
Anterior Primitive Streak
Blood progenitors 1
Blood progenitors 2
Cardiomyocytes
Caudal epiblast
Caudal Mesoderm
Caudal neurectoderm
Def. endoderm
Endothelium
Epiblast
Erythroid1
Erythroid2
Erythroid3
ExE ectoderm
ExE endoderm
ExE mesoderm
Forebrain/Midbrain/Hindbrain
Gut
Haematoendothelial progenitors
Intermediate mesoderm
Mesenchyme
Mixed mesoderm
Nascent mesoderm
Neural crest
NMP
Notochord
Paraxial mesoderm
Parietal endoderm
PGC
Pharyngeal mesoderm
Primitive Streak
Rostral neurectoderm
Somitic mesoderm
Spinal cord
Surface ectoderm
Visceral endoderm
D
Start from TE with high TE content (>=50%)
Derived from TE
Do not start from TE with high TE content (>=50%)
High TE percent
Start from TE with low TE content (<50%)
Start from TE
Do not start from TE with low TE content (<50%)
Other
TE
TE transcripts
(n = 137)
48 (35%)
28 (20.4%)
7 (5.1%)
54 (39.4%)
E
TE transcripts
(n = 137)
Number
100
50
0
Other
Overlap
NC genes

**Supplemental Figure S9 Evaluation of mouse scRNA-seq data from gastrulation to early organogenesis**

(A) Distribution of scRNA-seq reads in five non-overlapping genomic regions: protein-coding exons, TEs within the introns of protein-coding genes, other intronic regions of protein-coding genes, intergenic TEs, and other intergenic regions.

(B) Distribution of scRNA-seq reads in protein-coding exons, TE transcripts and other genomic regions.

(C) UMAP of scRNA-seq using the top 500 variable features that are protein-coding genes or TE transcripts.

(D) 82 of the 137 TE transcripts with strong tissue enrichment either initiate from TEs or have more than 50% of their exons composed of TEs.

(E) 47 of the 137 TE transcripts with strong tissue enrichment have been annotated by refSeq.

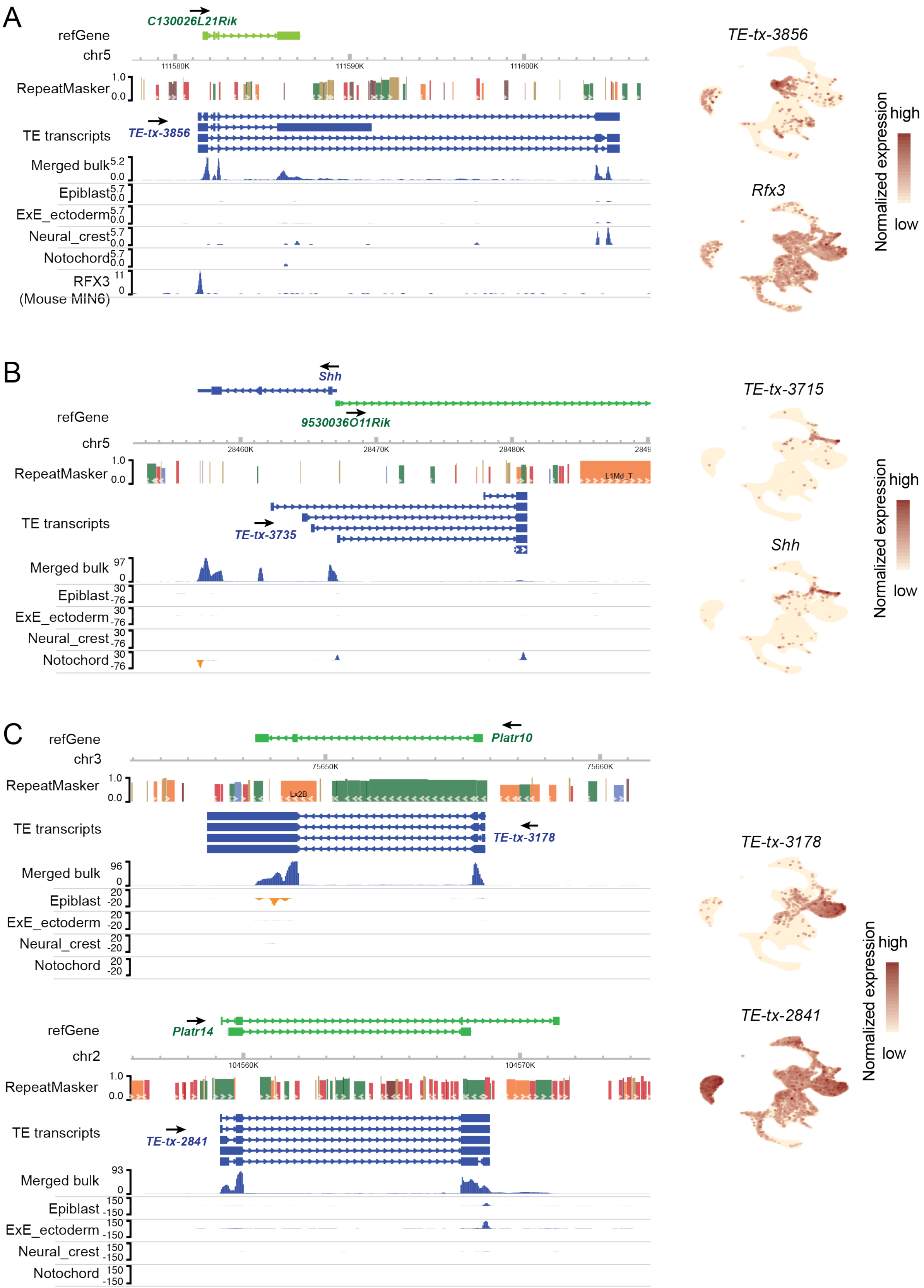
A
C130026L21Rik
refGene
chr5
111580K
111590K
111600K
RepeatMasker
TE transcripts
TE-tx-3856
Merged bulk
Epiblast
ExE_ectoderm
Neural_crest
Notochord
RFX3
(Mouse MIN6)
TE-tx-3856
Rfx3
Normalized expression
high
low
B
Shh
9530036O11Rik
refGene
chr5
28460K
28470K
28480K
RepeatMasker
L1Md_T
TE transcripts
TE-tx-3735
Merged bulk
Epiblast
ExE_ectoderm
Neural_crest
Notochord
TE-tx-3715
Shh
Normalized expression
high
low
C
refGene
Platr10
chr3
75650K
75660K
RepeatMasker
Lx2B
TE transcripts
TE-tx-3178
Merged bulk
Epiblast
ExE_ectoderm
Neural_crest
Notochord
refGene
Platr14
chr2
104560K
104570K
RepeatMasker
TE transcripts
TE-tx-2841
Merged bulk
Epiblast
ExE_ectoderm
Neural_crest
Notochord
TE-tx-3178
TE-tx-2841
Normalized expression
high
low

**Supplemental Figure S10 Examples of TE transcripts**

Left: genome browser view of assembled TE transcripts and scRNA-seq in selected tissues. Only Uniquely aligned reads were shown. RFX3 ChIP-seq track was obtained from Cistrome Data Browser (GEO: GSM1859216, CistromeDB: 56036). Right: Expression patterns of TE transcripts and related protein-coding genes.

**Supplemental Table S1 Summary of datasets utilized in this manuscript**

Descriptions and accession IDs of all the datasets utilized in this manuscript is listed in Supplemental_Table_S1.xlsx.

**Supplemental Table S2 Genomic locations of assembled TE transcripts**

Genomic locations of assembled 5299 TE transcripts are provided in Supplemental_Table_S2.xlsx.

**Supplemental Table S3 Expression patterns of 137 TE transcripts with strong tissue enrichment during mouse gastrulation and early organogenesis**

Expression patterns of 137 TE transcripts with strong tissue enrichment during mouse gastrulation and early organogenesis are summarized in Supplemental_Table_S3.xlsx.
